# A three-years assessment of *Ixodes ricinus*-borne pathogens in a French peri-urban forest

**DOI:** 10.1101/597013

**Authors:** Emilie Lejal, Maud Marsot, Karine Chalvet-Monfray, Jean-François Cosson, Sara Moutailler, Muriel Vayssier-Taussat, Thomas Pollet

## Abstract

**Background:** *Ixodes ricinus* is the predominant tick species in Europe and the primary pathogen vector for both humans and animals. These ticks are frequently involved in the transmission of *Borrelia burgdorferi* sensu lato, the causative agents of Lyme borreliosis. While much more is known about *I. ricinus* tick-borne pathogen composition, information about temporal tick-borne pathogen patterns remain scarce. These data are crucial for predicting seasonal/annual patterns which could improve understanding and prevent tick-borne diseases.

**Methods:** We examined tick-borne pathogen (TBPs) dynamics in *I. ricinus* collected monthly in a peri-urban forest over three consecutive years. In total, 998 nymphs were screened for 31 pathogenic species using high-throughput microfluidic real-time PCR.

**Results:** We detected DNA from *Anaplasma phagocytophilum* (5.3%), *Rickettsia helvetica* (4.5%), *Borrelia burgdorferi* s.l. (3.7%), *Borrelia miyamotoi* (1.2%), *Babesia venatorum* (1.5%) and *Rickettsia felis* (0.1%). Among all analysed ticks, 15.9% were infected by at least one of these microorganisms, and 1.3% were co-infected. Co-infections with *B. afzeli*/*B. garinii* and *B. garinii*/*B. spielmanii* were significantly over-represented. Moreover, significant variations in seasonal and/or inter-annual prevalence were observed for several pathogens (*R. helvetica, B. burgdorferi* s.l.*, B. miyamotoi*, and *A. phagocytophilum*).

**Conclusions:** Analysing TBPs prevalence in monthly sampled tick over three years allowed us to assess seasonal and inter-annual fluctuations of the prevalence of TBPs known to circulate in the sampled area, but also to punctually detect less common species. All these data emphasize that sporadic tick samplings are not sufficient to determine TBPs prevalence and that regular monitoring is necessary.

## Background

Ticks are obligatory hematophagous arthropods and consequently, are one of the most important pathogen vectors [1–3]. Lyme borreliosis (LB) is the most commonly reported tick-borne disease (TBD) in the northern hemisphere and is caused by bacteria belonging to the *Borrelia burgdorferi* s.l. complex. In Western Europe, *Ixodes ricinus* is known to be involved in the transmission of these bacteria to both humans and animals. This tick species has also been reported to be a vector for many other tick-borne pathogens (TBP) with potentially significant consequences for human and animal health (*Anaplasma*, *Rickettsia*, *Bartonella*, *Babesia*…) [4–9].

While multiple different pathogens have been identified and confirmed in *I. ricinus* ticks, very little is known about their seasonal and inter-annual variations. Time-series studies are thus crucial to understanding natural variability in microbial communities over time. Over the last decade, only a handful of surveys have assessed seasonal and monthly TBP variation patterns [10–14]. Although these results have heightened our general understanding of TBP dynamics, several of these studies were performed over short periods of less than two years, rendering it impossible to infer inter-annual discrepancies or to detect bias due to a particularly exceptional year. Only Coipan *et al*. [12] analysed several pathogenic genera in ticks sampled over more than two years. This study did demonstrate relationships between seasons and TBP prevalence (*Borrelia*, *Rickettsia*, *Anaplasma*, *Neoehrlichia*, and *Babesia*) in questing tick populations. These variations were mainly attributed to the varying availability of reservoir hosts.

Tick density is also heavily influenced by the presence of suitable hosts, most notably wild ungulates that sustain adults, thus enabling tick population renewal [15, 16]. However, it’s important to emphasise that immediate tick survival and questing activities are highly dependent on suitable and specific environmental conditions (temperatures between 8 to 24°C; and up to 80% humidity). Simultaneously, several studies have investigated whether pathogen presence influences tick behaviour. Herrmann and Gern [17, 18] suggested that *ricinus* infected with *B. burgdorferi* s.l. can tolerate increased levels of desiccation, and Neelakanta *et al*. [19] demonstrated that *I. scapularis* infected with *Anaplasma phagocytophilum* are more resistant to cold. The presence of these TBP could therefore enhance survival or questing activities of the infected ticks under challenging abiotic conditions, suggesting the existence of a potential link between pathogen prevalence in questing ticks and seasons.

Tick density and TBP prevalence can thus be influenced by several variables, and can therefore potentially fluctuate both seasonally and annually. Studying these dynamics is essential to better understanding and anticipating TBP risk.

Peri-urban forests containing both TBP-reservoir hosts and ticks, and which are highly frequented by people and their pets, represent a particularly interesting area to study tick and TBP dynamics. The Sénart forest, located to the south of Paris, harbours many large ungulates and abundant and diverse populations of other TBP reservoir hosts (bank voles, wood mice, Siberian chipmunks, roe deer, hedgehogs, …), and accommodates more than three million visitors every year. This forest is therefore particularly adapted to studying ticks and tick-borne pathogen dynamics.

In this study, we assessed the seasonal and inter-annual variability of *I. ricinus*-borne pathogens in the Sénart forest over three consecutive years (from April 2014 to May 2017), and determined whether any significant associations existed between these pathogens. We investigated a total of 31 pathogenic species (bacteria and parasites), belonging to 11 genera: *Borrelia*, *Anaplasma*, *Ehrlichia*, *Neoehrlichia* (only *Neoehrlichia mikurensis*), *Rickettsia*, *Bartonella*, *Francisella*, *Coxiella*, *Theileria*, *Babesia*, and *Hepatozoon*.

## Methods

### Tick collection

*I. ricinus*, nymphs and adults, were monthly collected during three years, from April 2014 to May 2017, in the Sénart forest in the south of Paris. Ticks were collected between 10 am and noon. Samplings were performed by dragging (Vassallo *et al*., 2000) on 10 transects of 10 square meters, localized on the parcel 96 (48°39′34.6″N 2°29′13.0″E, **Figure 1**). Dragging was always performed 3 consecutive times on each transect by the same persons to limit sampling bias. The presence of *Dermacentor* spp. was occasionally reported but no investigation has been led further. The presence of *I. ricinus* larvae was also sometimes noticed. Their small size making their individual DNA extraction and analysis difficult, we chose to not collect them. After morphological identification, ticks were stored at −80°C. In total 1167 ticks were collected.

**Figure 1.**
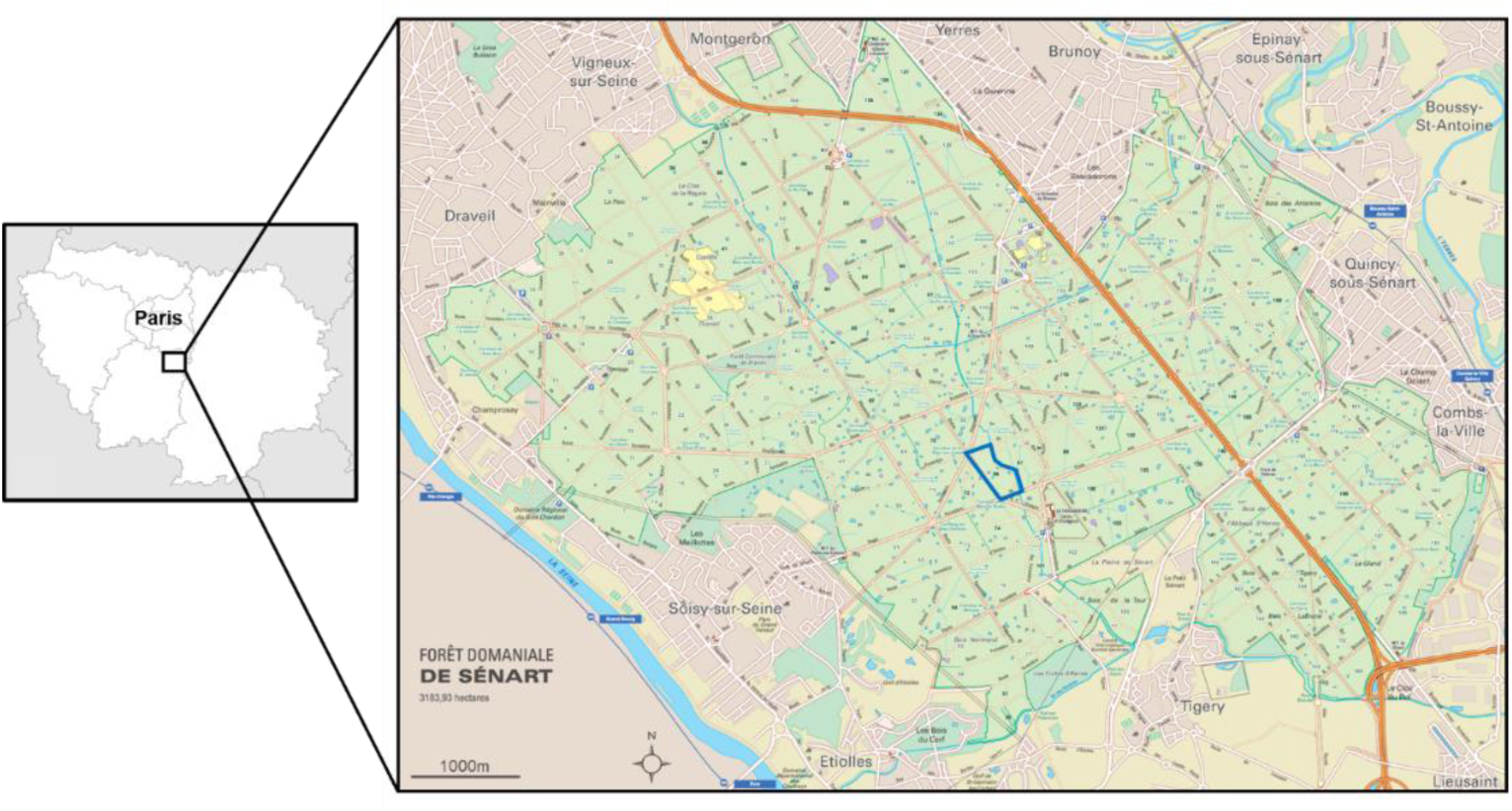
Sénart forest, location and parcel map. Sampling was made on the blue framed parcel.

### Tick washing, crushing and DNA extraction

Ticks were first washed once in ethanol 70% for 5 minutes and rinsed twice in sterile MilliQ water for 5 minutes. Ticks were then individually crushed in 375µL of Dulbecco’s Modified Eagle Medium (DMEM) with decomplemented Foetal Calf Serum (10%) and six steel beads using the homogenizer Precellys®24 Dual (Bertin, France) at 5500 rpm for 20 seconds.

DNA extractions were then performed on 100µL of tick crushing, using the DNA extraction kit NucleoSpin® Tissue (Macherey-Nagel, Germany), and following the standard protocol for human or animal tissue and cultured cells, from the step 2. DNA extracts were eluted in 50µL of elution buffer and then stored at −20°C until further use.

Two controls were performed: (1) the crushing control, corresponding to a DMEM tube in which crushing and DNA extraction were performed in the same conditions than on samples; and (2) the extraction control, corresponding to the DNA extraction step without tick samples.

### Tick-borne pathogens detection

A high-throughput screening of the most common bacterial and parasitic species known to circulate in ticks in Europe was performed. This allowed us to detect simultaneously the presence of 31 pathogenic species, 7 genera and 1 phylum: the Borrelia genus and eight Borrelia species (*B. burgdorferi* s.s., *B. afzelii*, *B. garinii*, *B. valaisiana*, *B. spielmanii*, *B. lusitaniae*, *B. bissettii* and *B. miyamotoi*); the Anaplasma genus and five Anaplasma species (*A. marginale*, *A. phagocytophilum*, *A. platys*, *A. centrale*, *A. bovis*); the Ehrlichia genus and *E. canis*; *Neoehrlichia mikurensis*; the Rickettsia genus and six Rickettsia species (*R. conorii, R. slovaca, R. massiliae, R. helvetica*, *R. aeshlimanii* and *R. felis*); the Bartonella genus and *B. henselae*; *Francisella tularensis*; *Coxiella burnettii*; the apicomplexa phylum and seven Babesia species (*B. divergens, B. microti, B. caballi, B. canis, B. venatorum, B. bovis, B. ovis)*, but also the two parasitic genus Theileria and Hepatozoon.

TBP DNA was detected using the BioMark™ real-time PCR system (Fluidigm, USA), a microfluidic system allowing to perform 48 or 96 real-time PCR reactions on 48 or 96 different samples as described in Michelet *et al*. [20] and Moutailler *et al*. [21]. Briefly, each sample and primers/probe set were included in individual wells. A pressure system allowed to load them on the chip, *via* microchannels, in individual reaction chambers of 10nL, where each sample will meet individually each primers/probe set.

### Primers and probes

Primers and probes used for this analysis have been developed and validated by Michelet *et al*. [20] and Gondard *et al*. [22]. They have been designed to specifically amplify DNA from pathogens (bacteria and parasites) which are usually found in ticks in Europe. Their sequences, amplicon size, as well as targeted genes and pathogens are presented in the Additional **table 1**. Please note that, due to potential cross-reactions between primer/probe combination (i.e. design) targeting *B. burgdorferi* s.s. and *B. spielmanii* with respectively *B. garinii*/*B. valaisiana* and *B. afzelii* DNA [described in 20], positive samples for the two formers were considered as negative when associated to the latter. Therefore, potential associations between *B. burgdorferi* s.s./*B. garinii*, *B. burgdorferi* s.s./*B. valaisiana* and *B. spielmanii*/*B. afzelii* cannot be detected and the co-infection percentage may be under-estimated.

**Table 1.**
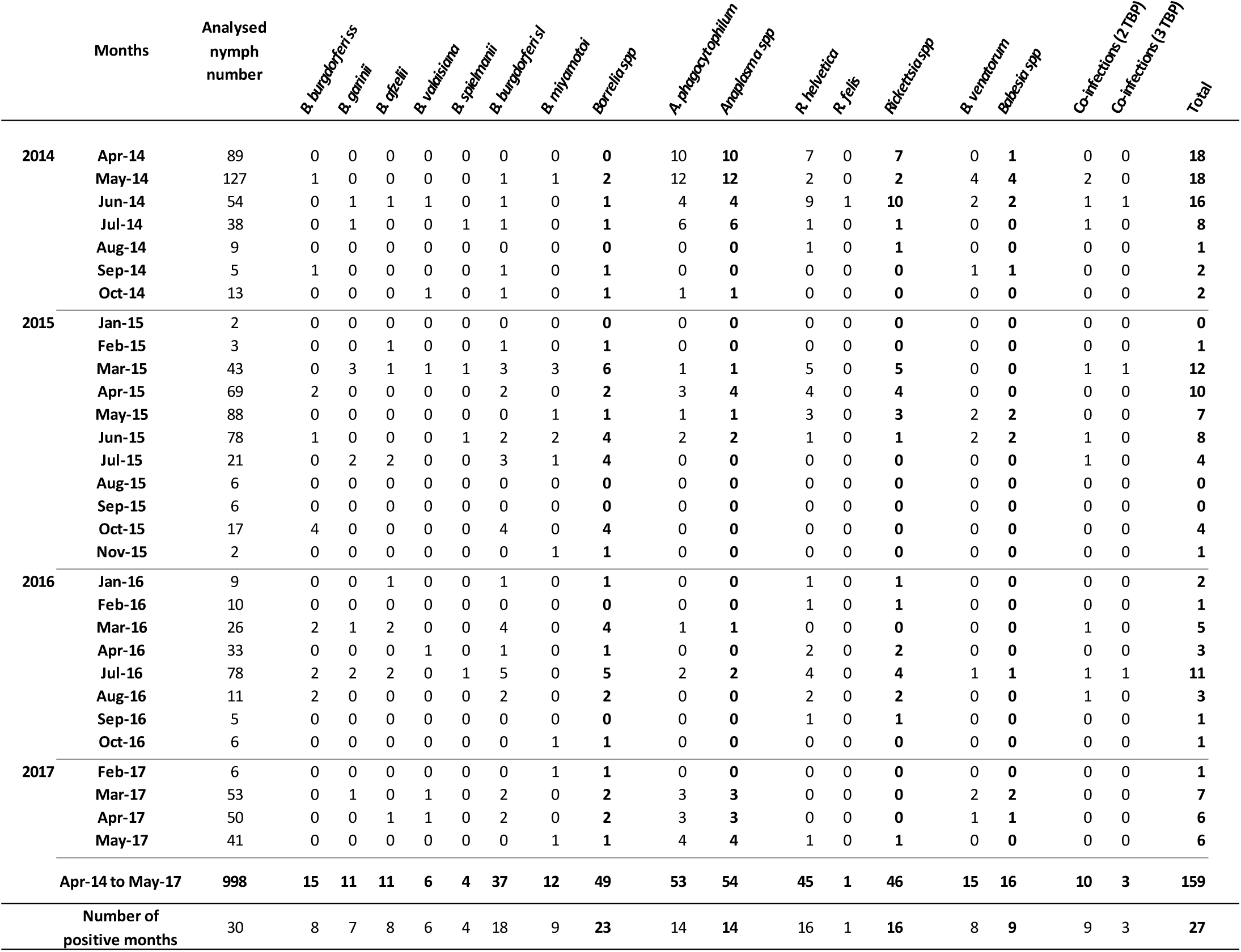
Summary table of the TBP detection study results.

### DNA pre-amplification

DNA pre-amplifiations were performed using the TaqMan PreAmp Master Mix (Applied Biosystems, France). Basically, the different primer pairs, used for the real time PCR, were pooled combining equal volume of primers with a final concentration of 0.2µM. For each sample, 1.25µL of DNA extract was pre-amplified using the Perfecta PreAmp SuperMix reagent (1x) and the 0.2x pool (0.05µM), in a final reactive volume of 5µL. PCR cycle comprised a first cycle at 98°C for 2 minutes, followed by 14 cycles with 2 steps, the first one at 95°C for 10 seconds and the second one at 60°C for 3 minutes. Pre-amplified DNA were then diluted (1:10) by addition of 45µL of sterile deionised water before use.

### High throughput real time PCR

For each pre-amplified sample, the BioMark™ real-time PCR system (Fluidigm, USA) was used for high-throughput microfluidic real-time PCR amplification using the 48.48 microfluidic dynamic array (Fluidigm Corporation, USA). Amplifications were performed using FAM- and black hole quencher (BHQ1)-labeled TaqMan probes with TaqMan Gene Expression Master Mix in accordance with manufacturer’s instructions (Applied Biosystems, France). Thermal cycling conditions were used as follows: 95°C for 5 min, 45 cycles at 95°C for 10 s, 60°C for 15 s, and 40°C for 10s. Data were acquired on the BioMark Real-Time PCR system and analysed using the Fluidigm Real-Time PCR Analysis software to obtain crossing point (CP) values. Three tick species control (*I. ricinus*, *Dermacentor reticulatus*, *Dermacentor marginatus*), one negative water control and one positive *Escherichia coli* control were included in each chip.

### Nested PCR and sequencing

Samples that were positive either only for species design but not for the genus design or only for the genus design and not for the species design were all re-analysed by nested PCR. We used primer pairs allowing to target another gene that the one tested into the fluidigm experiment. Their sequences, amplicon size, as well as targeted genes and pathogen genus are presented in the Additional **table 2**. Amplicons were then sequenced by the Eurofins company. Sequences were then analysed using the Bioedit software and compared to the database NCBI (National Center for Biotechnology Information) by sequence alignment using nucleotide BLAST (Basic Local Alignment Search Tool).

**Table 2.** Multivariable logistic regression models assessing the seasonal and yearly TBP prevalence variations in nymphs. Odds ratios and their associated 95% confidence intervals obtained from the best model of TBP seasonal and yearly prevalence in questing nymphs. REF = Reference – NA = Not Applicable.

| Model | TBP | Variable | Odds Ratio | 95% Confidence Interval |  |
| --- | --- | --- | --- | --- | --- |
|  |  |  |  | Low | High |
| (1) | *B. burgdorferi* s.l. | Spring |  | REF |  |
|  |  | Autumn | **4.53** | **1.50** | **12.49** * |
|  |  | Summer | 1.69 | 0.75 | 3.89 |
|  |  | Winter | 1.73 | 0.25 | 7.01 |
|  |  | 2014 |  | REF |  |
|  |  | 2015 | **2.93** | **1.12** | **9.14** * |
|  |  | 2016 | **4.48** | **1.60** | **14.53** * |
|  |  | 2017 | 2.45 | 0.57 | 9.95 |
| (2) | *B. miyamotoi* | Spring |  | REF |  |
|  |  | Autumn | 0.00 | NA | 8.3275E+218 |
|  |  | Summer | 0.00 | NA | 2.26397E+88 |
|  |  | Winter | **28.60** | **1.03** | **800.00** * |
| (3) | *A. phagocytophilum* | 2014 |  | REF |  |
|  |  | 2015 | **0.20** | **0.08** | **0.42** * |
|  |  | 2016 | **0.16** | **0.04** | **0.45** * |
|  |  | 2017 | 0.65 | 0.30 | 1.32 |
| (4) | *R. helvetica* | Spring |  | REF |  |
|  |  | Autumn | 0.00 | 0.00 | 6.7759E+11 |
|  |  | Summer | **3.10** | **1.27** | **7.85** * |
|  |  | Winter | 0.00 | NA | 1.1447E+145 |
|  |  | 2014 |  | REF |  |
|  |  | 2015 | 1.34 | 0.54 | 3.39 |
|  |  | 2016 | 0.81 | 0.12 | 3.24 |
|  |  | 2017 | **0.16** | **0.01** | **0.87** * |

### Statistical analysis

#### TBP prevalences at the seasonal and multi-annual scale

Differences in TBP prevalences were tested within and between years by using a multivariable logistic regression model. We considered the calendar season level for the within-year variability. Seasons were considered as following: Winter = December to February; Spring = March to May; Summer = June to August and Autumn = September to November.

Logistic regression models were developed using the TBP status of each nymph as the outcome measure and season, year and the interaction between season and year as explanatory variables. We performed four specific models for the following group/species of TBP: (1) *B. burgdorferi* s.l. (considering *B. burgdorferi* s.s., *B. garinii*, *B. afzelii*, *B. valaisiana* and *B. spielmanii*), (2) *B. miyamotoi*, (3) *A. phagocytophilum*, and (4) *R. helvetica*. The models were constructed from a generalized linear model [GLM, 23] using a binomial distribution (logit link). Model assessment was based on Akaike information criterion (AIC). Results were expressed as odds ratios (OR) and 95% confidence intervals. Statistical computations were performed in R 3.5.1. [24].

#### Statistical modelling of tick-borne pathogen associations

We tested the associations between the TBP species that belonged to the co-infection profiles of nymphs found in this study. We used the association screening approach [25]. For a given number of pathogen species tested (NP), the number of possible combination (NC) was calculated as NC = 2^NP^. Assuming similar pathogen prevalence as those observed, a simulated dataset was built as an absence/presence matrix with hosts in lines and pathogen combinations in columns. With 5 000 simulations, we obtained the NC statistic distributions. We estimated a 95% confidence interval to obtain a profile that includes simultaneously all the combinations. From this profile, we inferred for each combination two quantiles, *Qinf* and *Qsup*. A global test was based on the 95% confidence envelope. When H0 was rejected, the local tests were based on the NC confidence intervals: [*Qinf*; *Qsup*][25].

## Results

### Tick temporal dynamics

From April 2014 to May 2017, a total of 1167 *Ixodes ricinus* ticks were collected in the Sénart forest in the south of Paris (**Figure 1**). Please note that May and June 2016 were unfortunately not sampled due to logistic issues. Collected ticks were composed of 1098 nymphs, 35 females and 34 males. Adults were sporadically detected all over the three years Due to their low total abundance (more than 10 fold less compared with nymphs), we decided to focus our temporal analysis on nymphs. The temporal dynamics of nymph densities over the three years is shown in **Figure 2**. Nymph densities followed similar patterns from one year to another, with a main peak of activity observed every year in spring, a strong decrease during summer and a second peak, smaller, observed in Autumn (**Figure 2**). In January and February, the average densities were less than 10 questing nymphs/100m^2^. A clear rise was observed from March to May reaching an average peak of 95 nymphs/100m^2^ in May. We then observed a decrease in summer up to a minimum average of 5 nymphs/100m^2^ in September. The nymph densities increased slightly in October to reach an average of 13 nymphs/100m^2^, before finally decreasing in November (2 nymphs/100m^2^, sampled in 2015).

**Figure 2.**
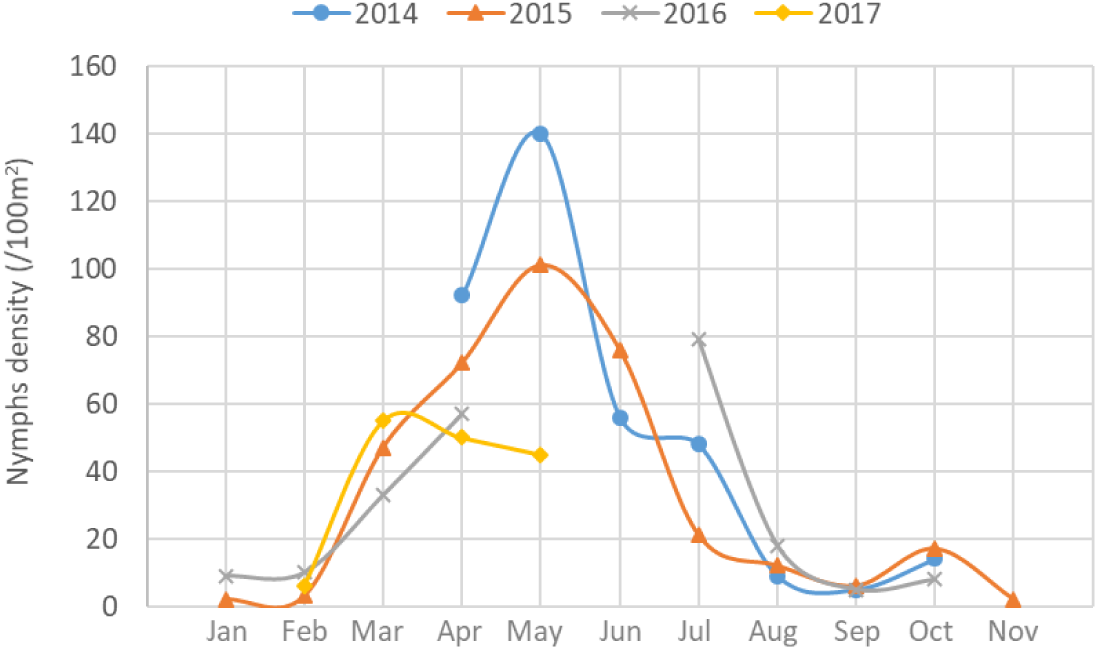
*Ixodes ricinus* nymphs monthly density (/100m^2^) in 2014, 2015, 2016 and 2017. Ticks were sampled from April 2014 to May 2017. Please note that May and June 2016 were unfortunately not sampled.

### Detected pathogens and their prevalence in tick population

Due to technical problems, DNA was extracted and analysed only from 1044 nymphs among the 1098 previously mentioned. 46 were negative for at least one positive control and thus have been removed from the analysis. From the 998 remaining DNA samples, 15.9% [13.7%, 18.3%] were positive for at least one tested pathogen, which belong to three bacterial and one protozoan genera: *Anaplasma*, *Borrelia*, *Rickettsia* and *Babesia* (**Table 1**).

Pathogens belonging to the *Anaplasma* genus were detected in 5.4% [4.1%, 7.0%] of collected ticks. Most of them were positive for *Anaplasma phagocytophilum* (5.3% of all the samples) and one DNA sample was only positive for the primers/probe combination specific to *Anaplasma* spp.. This sample was confirmed by nested PCR and the amplicon was then sequenced. The BLAST analysis on NCBI showed that this sequence matched at 99% of identity with four different *Anaplasma* species (*A. phagocytophilum*, *A. marginale*, *A. ovis* and *A. centrale*). Therefore, this sample was only considered as positive for *Anaplasma* spp..

Two species of *Rickettsia* were detected in questing *I. ricinus* nymphs. *Rickettsia helvetica* was the most prevalent and was detected in 4.5% [3.3%, 6.0%] of nymphs. *Rickettsia felis* was detected in only one nymph (0.1% [.003%, 0.6%]). The presence of *R. felis* DNA was confirmed by nested PCR and sequencing of the *ompB* gene. The obtained sequence (GenBank accession number MN267050) matched with the corresponding gene sequence of *R. felis* (GenBank accession numbers GU182892.1) with 100% of identity and 100% of query cover.

The genus *Borrelia* was represented by six different species detected in 4.9% [3.7%, 6.4%] of the surveyed nymphs. Five belonged to the LB group (3.7% [3.7%, 6.4%]), including *B. burgdorferi* s.s. (1.5% [0.8%, 2.5%]), *B. garinii* (1.1% [0.6%, 2.0%]), *B. afzelii* (1.1% [0.6%, 2.0%]), *B. valaisiana* (0.6% [0.2%, 1.3%]) and *B. spielmanii* (0.4% [0.1%, 1.0%]). DNA of *Borrelia miyamotoi*, belonging to the relapsing fever group, was detected in 1.2% [0.6%, 2.1%] of the collected nymphs.

DNA from two species of *Babesia* were detected in questing nymph with the microfluidic PCR: *Babesia venatorum* (1.5% [0.8%, 2.5%] of ticks) and *Babesia divergens* (0.1% [0.003%, 0.6%]), detected in one tick). A deeper investigation of the *B. divergens* positive sample, by nested PCR on the 18s rRNA gene and amplicon sequencing (GenBank accession number MN296295), allowed us to finally identify the DNA of *B. capreoli*, closely related species to *B. divergens*, circulating in wild ruminants and unable to infect Human and cattle erythrocytes [26].

### Temporal patterns of TBP in *Ixodes ricinus* nymphs

#### TBP prevalence at the monthly scale

In the following paragraph and corresponding figures, prevalences are those obtained for months with at least nine collected ticks.

Global infection rates fluctuated from 8% [3.3%, 15.7%] in May 2015 to 29.6% [18.0%, 43.6%] in June 2014 (**Figure 3**). At the genus level, variations in TBP prevalences and the number of months for which at least one tick was positive for each tested TBP are presented in **Figure 4** and **Table 1**.

**Figure 3.**
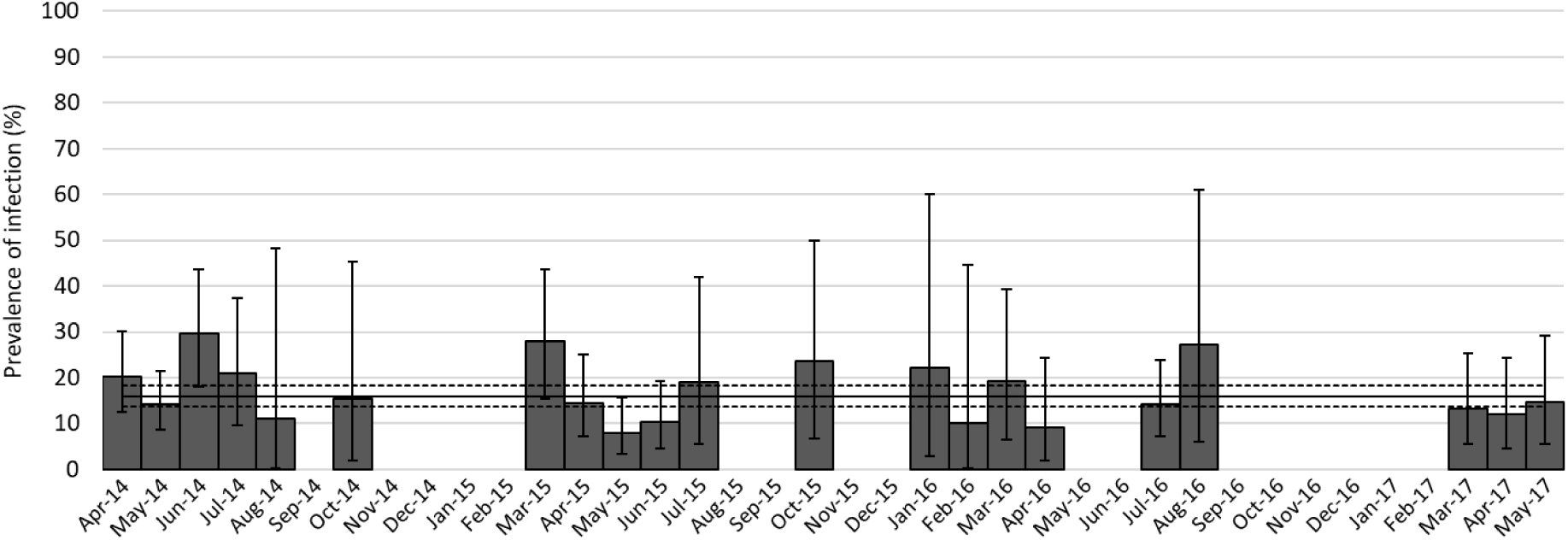
Nymph infection rate per month for at least one tested pathogen. Months with less than nine nymphs sampled have not been considered for percentage calculation. Error bars represent confidence intervals of the percentage.

**Figure 4.**
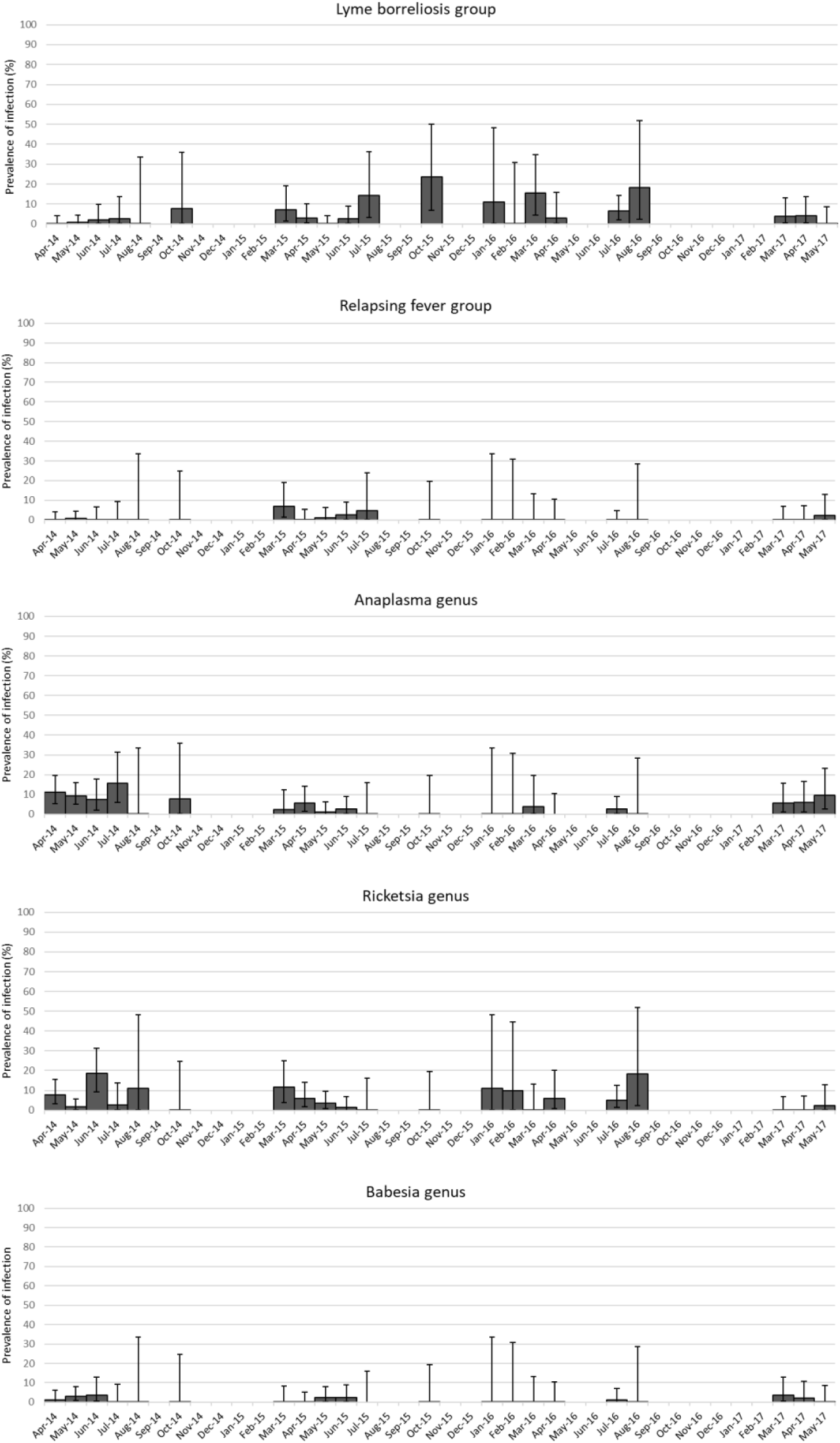
Nymph infection rate and confidence intervals per month for the different TBP. Months with less than nine nymphs sampled have not been considered. Error bars represent confidence intervals of the percentage.

DNA from pathogens belonging to both genera *Rickettsia* and *Anaplasma* were detected respectively in 16 and 14 of the 30 sampled months respectively. When detected, prevalences fluctuated from 1.3% [0.03%, 6.9%] (June 2015) to 18.5% [9.3%, 31.4%] (June *A.* 2014) for *Rickettsia* and from 1.1% [0.03%, 6.2%] (May 2015) to 15.8% [6.0%, 31.3%] (July 2014) for *Anaplasma*. Both genera are mainly represented by one species: *R. helvetica* and *A. phagocytophilum* that are the most frequently detected species (16 and 14 /30 months respectively). These two species were found each year.

DNA from members of the *Borrelia* genus was detected in 23 of the 30 sampled months. This bacterial genus displayed the highest variability with prevalences fluctuating from 1.1% [0.03%, 6.2%] (May 2015) to 23.5% [6.8%, 49.9%] (October 2015). DNA from members of the LB group was detected in 18 of the 30 sampled months with prevalences ranging from 0.8% [0.03%, 6.2%] in May 2014 to 23.5% [6.8%, 49.9%] in October 2015. The most frequently identified species were *B. burgdorferi* s.s. (8 / 30 sampled months), *B. afzelii* (8 / 30) and *B. garinii* (7 / 30). DNA from these species was regularly detected over the three studied years. Conversely, *B. valaisiana* (6 / 30) and *B. spielmanii* (4 / 30) DNA was not detected during 11 (from April 2015 to March 2016) and 9 (from July 2015 to April 2016) consecutive months respectively. *Borrelia miyamotoi* (relapsing fever group) DNA was detected 9 times over the 30 sampled months with prevalences ranging from 0.8% [0.02%, 4.3%] in May 2014 to 7% [1.5%, 19.1%] in March 2015.

For parasites, DNA from the genus *Babesia* was detected in 9 months out of 30 sampled months. Prevalences presented the lowest variability ranging from 1.1% [0.03%, 6.1%] in April 2014 to 3.8% [0.5%, 13.0%] in March 2017 (**Figure 4**). The main detected species was *venatorum* that was detected 9 times over 30 samplings and not detected during 9 consecutive sampled months, from June 2015 to April 2016.

#### TBP prevalences at the seasonal and multi-annual scale

In order to determine if the prevalence of TBP was different within and between years, a multivariable logistic regression model was performed. The spring season and the year 2014 have been considered as references for the seasonal and yearly effect respectively. Because some TBPs had too low prevalences in the nymph population (producing unreliable statistics), analyses were only performed on the most prevalent TBPs: *A. phagocytophilum*, *R. helvetica*, *B. burgdorferi* s.l., *B. miyamotoi* and *B. venatorum*.

Significant differences were observed at the seasonal scale (**Figure 5**, **Table 2**) for *R. helvetica* (higher in summer compared to spring), *B. burgdorferi* s.l. (higher in autumn compared to spring) and *B. miyamotoi* (higher in winter than in spring). Please note that the smallest number of sampled ticks (30 in total) was found in winter, and that the difference observed for *B. miyamotoi* in winter corresponded to only one infected tick collected in February 2017.

**Figure 5.**
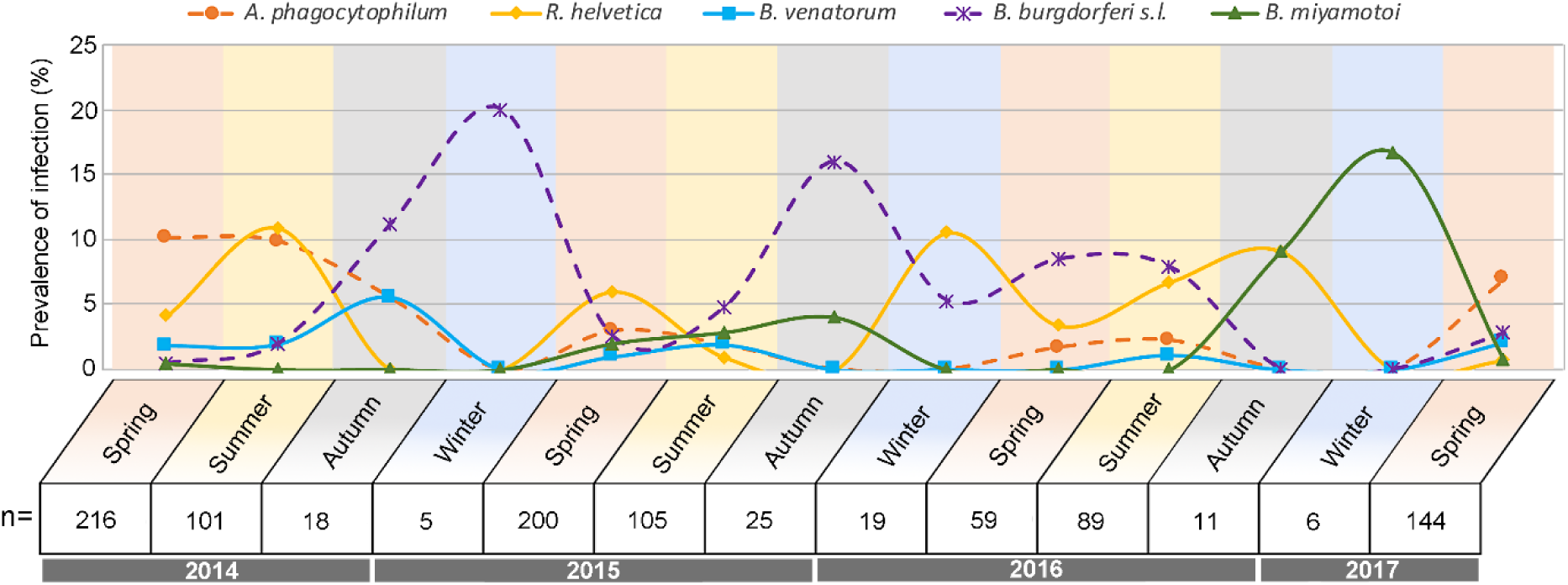
Percentage of positive nymphs per season for the most prevalent TBPs. Winter (pastel blue background) = January to February – Spring (pastel orange background) = March to May – Summer (pastel yellow background) = June to August – Autumn (light grey background) = September to November – n = Number of analysed ticks.

Significant differences were also observed between years for bacteria belonging to the complex *B. burgdorferi* s.l. with higher infection rates in 2015 and 2016 compared to 2014; for *A. phagocytophilum*, which was lower in 2015 and in 2016 compared to 2014 and for *R. helvetica*, which was lower in 2017 than in 2014. However, please note that samplings were only performed from January to May in 2017. No significant differences were observed according to season or year for *B. venatorum*.

#### Pathogen associations

Among all the sampled ticks, 1% [0.5%, 1.8%] were co-infected with two pathogens and 0.3% [0.006%, 0.8%] were co-infected with three pathogens. Eight different co-infection profiles were found (**Table 3**). In most of cases (7/13), these co-infections concerned species belonging to the *Borrelia* genus: *B. garinii*/*B. afzelii*; *B. garinii*/*B. spielmanii*; *B. garinii*/*B. afzelii*/*B. valaisiana* and *B. garinii*/*B. valaisiana*/*B. spielmanii*. Co-infections profiles with species belonging to different genus were also observed: *A. phagocytophilum*/*B. venatorum*; *A. phagocytophilum*/*R. helvetica*; *B. burgdorferi* s.s./*R. helvetica* and *B. garinii*/*B. afzelii*/*R. helvetica*. All these associations between pathogens were tested using the association screening approach (Vaumourin *et al*., 2014). Compared to a random analysis, no associations were found to be under represented while two were over represented: the first one between *B. garinii* and *B. afzelii* (observation = 3; min expected = 0; max expected = 2), and the second one between *B. garinii* and *B. spielmanii* (observation = 2; min expected = 0; max expected = 1).

**Table 3.** Summary table of the reported co-infection profiles.

| *B. burgdorferi s.s.* | *B. garinii* | *B. afzelii* | *B. valaisiana* | *B. spielmanii* | *A. phagocytophilum* | *R. helvetica* | *B. venatorum* | Co-occurrences number |
| --- | --- | --- | --- | --- | --- | --- | --- | --- |
|  | X | X |  |  |  |  |  | 3 |
|  | X |  |  | X |  |  |  | 2 |
|  | X | X | X |  |  |  |  | 1 |
|  | X |  | X | X |  |  |  | 1 |
|  |  |  |  |  | X |  | X | 3 |
|  |  |  |  |  | X | X |  | 1 |
| X |  |  |  |  |  | X |  | 1 |
|  | X | X |  |  |  | X |  | 1 |

## Discussion

### *Ixodes ricinus* density and seasonal dynamics

This three-year survey demonstrated a clear seasonal pattern in *I. ricinus* density, with a marked peak of questing nymphs in spring and a smaller peak in autumn. Low, but present activity was detected in winter, as has been observed in Germany [27]. In addition to these general patterns, some unexpected data were observed, the most striking being no peak activity in spring 2017 (April and May) with tick densities very similar to those recorded in March. Abiotic factors such as temperature, relative humidity, and rainfall, or fluctuating host numbers in the sampling area are known to influence questing tick abundance and activity patterns [28–33] and could explain these unusual observations. It’s important to note that 2017 was distinguished by an abnormally wet March, with total rainfall much higher than that recorded in previous years in the same area (71.3, compared to 11.2, 33.6, and 61.7 mm rain/month in 2014, 2015, and 2016, respectively). Interestingly, the increased March rainfall was followed by an April drought (7.9 mm of rain/month in 2017, compared to 48.4, 27.2, and 66.2 mm rain/month in 2014, 2015, and 2016, respectively) (rainfall data estimated from the Orly station, Metéo-France data; https://donneespubliques.meteofrance.fr/?fond=produit&id_produit=90&id_rubrique=32). These unusual meteorological characteristics could explain the stable tick density from March to May 2017. Thereby, this finding clearly shows that the bimodal tick activity pattern usually observed during this study can punctually change with exceptional environmental conditions, reinforcing the importance of regular monitoring.

### *Ixodes ricinus*-borne pathogen composition and prevalence over the three years

Most of the detected pathogen species corresponded to micro-organisms known to circulate in the Western Palearctic [34–41]. However, several species belonging to the *Bartonella* and *Francisella* genera, previously reported in the studied area [39, 42], were not detected. The most prevalent pathogen species were *A. phagocytophilum* (5.4% of the examined nymphs), *R. helvetica* (4.5%), and *B. burgdorferi* s.l. (3.7%). Both high- and low-prevalence TBP were consistently detected in the sampling area for the duration of the study. Although prevalences varied between different TBPs, and some were not detected for long periods, they were all detected recurrently. Continued detection is consistent with the year-round presence of reservoir hosts in the sampling area (wood mice, bank voles, Siberian chipmunks, roe deer, common blackbird, European robin, song thrush…) [33,43,44]. The continued presence of reservoir hosts could facilitate the circulation of dominant species, and maintain, even at low rates, less prevalent pathogen species that may not be detected by a single sampling. This does support to regularly studying TBP temporal dynamics, to improve the assessment of their prevalence.

We also detected in a single tick, the DNA of the emergent human pathogen *R. felis*. Its detection is particularly scheming as this bacteria is known to be mainly transmitted from cat to cat *via* fleas, with human contamination arising from cat or flea bites. As we only detected DNA from *R. felis*, we cannot exclude that this detection could correspond to remnant DNA from the previous blood meal. Nevertheless, several studies have already detected the presence of *R. felis* or *R. felis*-like organisms in hematophagous arthropods [see in 45,46], including in ticks collected from natural environments [47], and notably in two studies performed on questing *I. ricinus*, including one based on RNA detection [48, 49]. Rarely investigated in studies dealing with TBP, the repeated detection of *R. felis* should encourage increased surveillance for this spotted fever-causing pathogen in humans. Let’s finally note that all these findings suggest that a punctual sampling would certainly not facilitate the detection of this pathogen, again highlighting the importance of collecting and analysing ticks at a large temporal scale.

*B. divergens* and *B. capreoli* are two closely related species. During this study, we found out that the design initially used to detect *B. divergens* was actually also able to detect the DNA of *B. capreoli*. While *B. divergens* is responsible of babesiosis in human and cattle, *B. capreoli* is only able to colonize erythrocytes from deer, therefore presenting no threat for human or livestock [26]. These results emphases the importance of confirmation and careful interpretations of microfluidic real time PCR [50].

### Seasonal and inter-annual dynamics of *I. ricinus*-borne pathogens

Improving the prevention of TBD requires a better understanding of their temporal— and in particular—their seasonal dynamics. However, only a few studies have addressed these issues during a minimum three-year period [12, 13]. As ticks were collected monthly for over three years in this study, we detected significant seasonal or annual infection rate fluctuations for four TBP: *R. helvetica*, *B. burgdorferi* s.l., *B. miyamotoi*, and *A. phagocytophilum*. Note that the statistically significant highest prevalence of *B. miyamotoi* in winter is only due to the detection of one positive tick sampled during winter in 2017. In our opinion, this result alone is insufficient to presume that *B. miyamotoi* have an increased winter prevalence. However, we can observe that even if very few ticks are questing during these periods, they may carry TBP.

While significant seasonal and annual differences were observed for *B. miyamotoi* and *A. phagocytophilum*, respectively, the presence of *R. helvetica* and *B. burgdorferi* s.l. varied significantly according to both seasons and years. None of these micro-organisms presented a similar pattern to any others. Comparing our results to the pluri-annual studies previously mentioned, we observe that only *R. helvetica* presented similar seasonal patterns [12]. This finding again emphasises how the season, the year or the sampling area can influence TBP presence and prevalence in questing tick populations.

The most common explanation for temporal variations in TBP prevalences is the variable availability of reservoir hosts during tick previous stage feeding. This hypothesis was already suggested by Coipan *et al.* [12] while they observed that several micro-organisms, assumed to share the same reservoir host, also presented similar seasonal patterns. Because the tick lifecycle is fundamentally linked to its host, any changes to the available host spectrum will undoubtedly influence TBP prevalence in the tick community [51]. However, because the entire tick life cycle is pluri-annual, it is difficult anyway to know if nymphs questing at a same time did perform their previous blood meal at the same period. The same generation of questing nymphs could come from larvae that would have fed at different moment and thus potentially upon different host species. An alternative hypothesis, based on both the presence of pathogens and the tick physiology, could also explain these patterns. Carrying certain TBP was shown to improve tick resistance to challenging abiotic conditions. Herrmann and Gern [17, 18] demonstrated that ticks carrying *Borrelia* species exhibited higher survival rates in desiccating conditions and a lower tendency to move to favourable conditions for maintaining water balance than non-infected ticks. This was associated to a higher reserve of energy in Borrelia infected ticks [52] which would therefore exhibit higher resistance capacities to hydric stress notably. This hypothesis would explain the higher prevalence of *Borrelia*-infected questing ticks observed during or after the summer period in the present study and in those of Coipan and Takken [12, 13]. Similarly, Neelakanta [19] demonstrated a higher expression of *iafgp* gene, coding for an antifreeze glycoprotein, in *A. phagocytophilum*-infected ticks. This thus conferred to ticks a stronger resistance to cold that could lead to higher prevalence of *A. phagocytophilum*-infected questing ticks during or just after winter. This hypothesis was not consistent with our data as *A. phagocytophilum* was not observed in greater prevalence during the cold seasons of our study.

Our results, in combination with those from the literature, support the hypothesis that TBP prevalence is influenced by both biotic and abiotic factors, and suggest one more time that sporadic samplings are insufficient to assess it.

### Pathogen co-occurrence

Tick co-infections are being identified more and more frequently [21,41,49,53–57]. Clinical co-infections with several TBP are commonly reported [58–60] and are known to affect both disease symptoms and severity [61, 62]. It is thus essential to investigate TBP associations in ticks, to better identify potential clinical co-infections and to improve epidemiological knowledge of TBD.

In this longitudinal three-year study, two TBP associations were significantly over-represented compared to a random distribution: the first one was between *B. garinii* and *B. afzelii*, as has been previously observed in studies using similar detection tools [21, 41], or different methods [16s rRNA gene sequencing, 63]; the second one was between *B. garinii* and *B. spielmanii*. Interestingly, these findings contrast with published results on *Ixodes ricinus* TBP. While performing a meta-analysis on data published from 2010 to 2016, Strnad *et al* [64] observed a negative correlation between *B. garinii* and *B. afzelii*. Similarly, Herrmann *et al.* [65] also detected a negative co-occurrence between these two species following the analysis of 7400 nymphs collected over three years. These results are coherent considering the host specificity of these Borrelia species. Indeed, *B. garinii* doesn’t share the same reservoir host (birds) than *B. afzelii* or *B. spielmanii* (wood mice and bank voles, or hazel and garden dormice) [66–71], and none of these species are known to be transmitted transovarially.

Even though the associations we identified were statistically “over-represented”, in actual fact we only observed one more association than the fixed over-representation threshold (i.e. observed associations = 3 and 2; minimum expected = 0 and 0; maximum expected = 2 and 1; for *B. garinii*/*B. afzelii* and *B. garinii*/*B. spielmanii* associations, respectively). This indicates that caution should be applied when drawing conclusions about permanent associations between these different bacteria in ticks. Several different hypotheses could potentially explain these associations in the same nymph. Firstly, hosts are likely to carry several adjacent feeding ticks. This phenomenon, known as co-feeding, could promote pathogen exchange between ticks even in the absence of systemic host infection [72]. Secondly, as discussed by van Duijvendijk *et al*. [73], when bloodmeals are disrupted due to host grooming, immune response or death, ticks may feed on more than one host to completely engorge, and consequently be exposed to several pathogens. Thirdly, despite these TBP species segregating between bird and rodent hosts, all of them have been detected in hedgehogs [74, 75], and *B. afzelii* and *B. garinii* have been simultaneously detected in one Siberian chipmunk [44]. Both of these mammals were found to host a large number of tick larvae [44, 76], and Siberian chipmunks have been reported to induce higher *B. burgdorferi* s.l. infection rates in nymphs, compared to bank voles and wood mice [44] in the Sénart forest. A last hypothesis might be that our analyses methods are unable to distinguish the rodent-circulating *B. garinii* OspA serotype 4 (corresponding to *B. bavariensis*) [77] from other *B. garinii* serotypes.

Associations between *B. garinii* and *B. valaisiana* are frequently reported, which isn’t surprising as these species share the same reservoir host [78]. This association was the most common TBP association in a meta-analysis of literature published between 1984 and 2003 [79], and has been reported several times since in later studies [11,65,80]. While we observed this association twice, both times in association with a third *Borrelia* species, either *B. afzelii* or *B. spielmanii*, it was not significantly over-represented compared to a random distribution. Among the three previously mentioned studies, only Herrmann *et al.* [65] demonstrated that this association was over-represented when compared to a randomly sampled analysis. However, our study was performed on a much smaller dataset (998 versus 7400 analysed nymphs), with a halved co-infection percentage (1.3% versus 3%), indicating that our statistical analysis may be less powerful, which could explain why this association wasn’t detected.

These contrasting tick pathogen association results highlight the complexity in clearly identifying pathogen associations in field-collected ticks. Several other parameters can also potentially influence pathogen association (host spectrum within the studied area, sample size influencing analytical statistical power, identification bias…). In this context, performing investigations under controlled conditions (suitable TBP growing and tick breeding systems…) will be a crucial future step to experimentally test these different associations and improve our knowledge on TBP co-occurrence.

## Conclusions

This three-year study of *I. ricinus-*borne pathogens; (1) identified several TBP previously reported in the area, consistent with reservoir host availability; (2) allowed the surprising detection of *R. felis* DNA, a micro-organism rarely reported in questing ticks; (3) highlighted significant variations in seasonal and inter-annual pathogen prevalence; and finally (4) identified several unexpected co-occurrences between pathogens belonging to the *B. burgdorferi* s.l. complex. All these results represent another step towards understanding the TBPs ecology and emphase the need to perform longitudinal study. Especially since the main factors that are supposed to influence tick and TBP ecology may change in the next coming years with climate changes. Associated to other factors such as host information or meteorological measures, this kind of data is crucial to allow a better understanding of TBP ecology and TBD epidemiology.

## Additional files

**Additional file 1: Table S1.** Targeted gene, amplicon size, primers and probe sequences used for TBP and Tick species detection.

**Additional file 2: Table S2.** Targeted gene and primers sequences used for results confirmation.

## Abbreviation

TBP: tick-borne pathogens; TBD: tick-borne disease; LB: lyme borreliosis; *I. ricinus*: *Ixodes ricinus*; s.l.: sensu lato; s.s.: sensu stricto; *B. burgdorferi*: *Borrelia burgdorferi*; *B. afzelii*: *Borrelia afzelii*; *B. garinii*: *Borrelia garinii*; *B. valaisiana*: *Borrelia valaisiana*; *B. spielmanii*: *Borrelia spielmanii*; *B. lusitaniae*: *Borrelia lusitaniae*; *B. bissettii*: *Borrelia bissettii*; *B. miyamotoi*: *Borrelia miyamotoi*; *A. phagocytophilum*: *Anaplasma phagocytophilum*; *A. marginale*: *Anaplasma marginale*; *A. platys*: *Anaplasma platys*; *A. centrale*: *Anaplasma centrale*; *A. bovis*: *Anaplasma bovis*; *B. venatorum*: *Babesia venatorum*; *B. divergens*: *Babesia divergens*; *B. capreoli*: *Babesia capreoli*; *B. microti*: *Babesia microti*; *B. caballi*: *Babesia caballi*; *B. canis*: *Babesia canis*; *B. bovis*: *Babesia bovis*; *B. ovis*: *Babesia ovis*; *B. henselae*: *Bartonella henselae*; *E. canis*: *Ehrlichia* canis; *R. helvetica*: *Rickettsia helvetica*; *R. felis*: *Rickettsia felis*; *R. conorii*: *Rickettsia conorii*; *R. slovaca*: *Rickettsia slovaca*; *R. massiliae*: *Rickettsia massiliae*; *aeshlimanii*: *Rickettsia aeshlimanii*; DMEM: Dulbecco’s Modified Eagle Medium; NCBI: National Center for Biotechnology Information; BLAST: Basic Local Alignment Search Tool; GLM: Generalized Linear Model; AIC: Akaike Information Criterion; OR: Odds Ratios; NP: Number of Pathogen species tested; NC: Number of possible Combination; iafgp: *Ixodes scapularis* antifreeze glycoprotein

## Acknowledgments

The authors would like to thanks the financers of this project, that are the French National Institute of Agronomic Research (INRA): the métaprogramme “Metaomics and microbial ecosystems” (MEM) and the Métaprogramme “Adaptation of Agriculture and Forests to Climate Change” (ACCAF) that funded the tick collection from the CC-EID project. We also would like to thank the Ile-de-France region that funded the salary of Emilie Lejal, the PhD student working on this project. We are very grateful to all the persons that brought their help to the support of the technical part of this project: Julie Cat, Valérie Poux and members of the VECTOTIQ team for their involvement in tick samplings and identification; Elodie Devillers, for her precious help to learn the Microfluidic PCR approach and also Sabine Delannoy and the IdentyPath genomic platform that allowed us to perform the tick-borne pathogens screening. Thank you as well to Ladislav Šimo for his precious help in figure improvement.

## Authors’ contributions

Conceived and designed the experiments: TP, MVT, JFC, KCM, EL. Performed the experiments: EL. Analysed the data: EL, TP, SM, MM, KCM. Wrote the paper: EL, MM, KCM, JFC, SM, MVT, TP

---

**Table S1.**
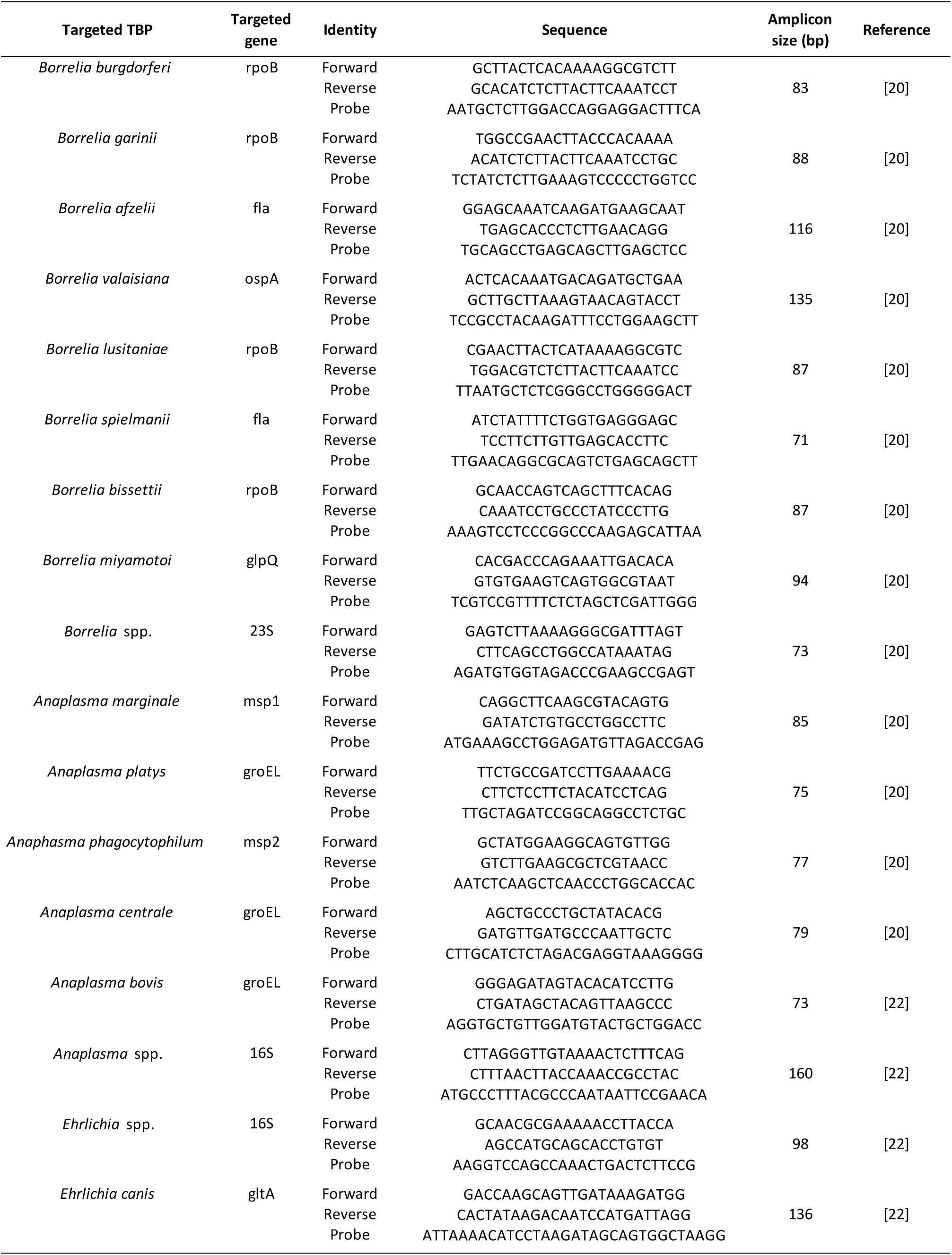
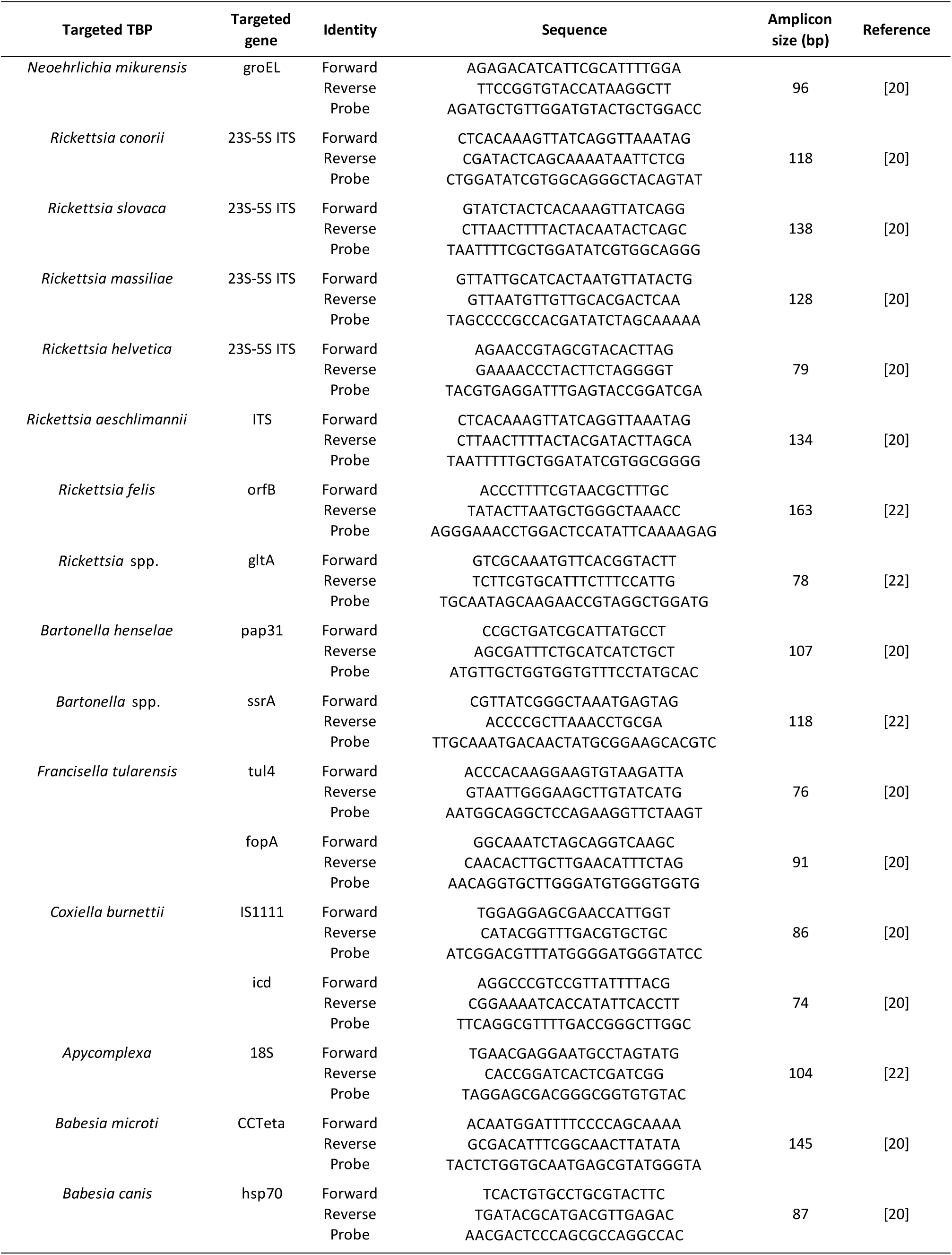
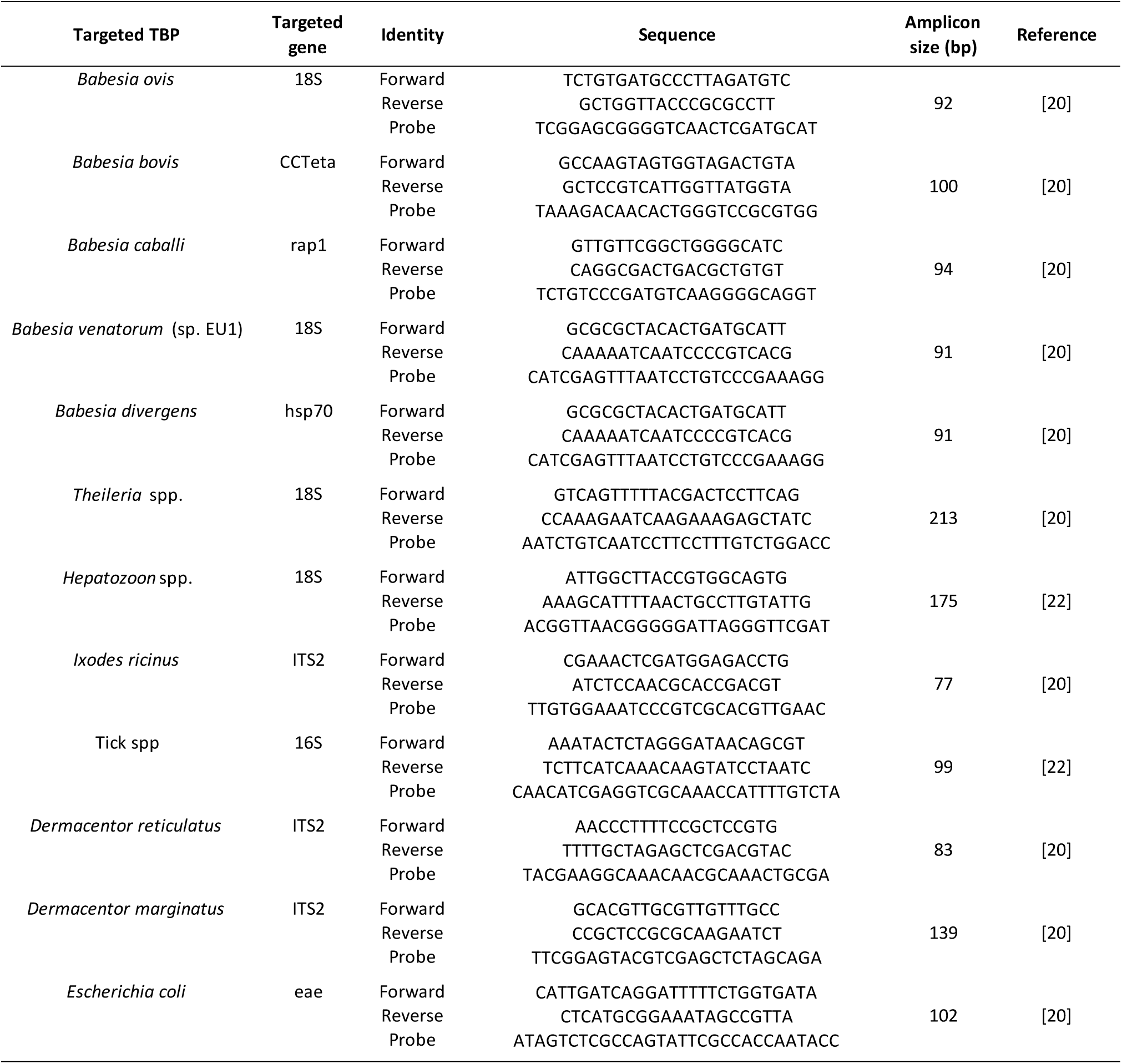

**Table S2.**
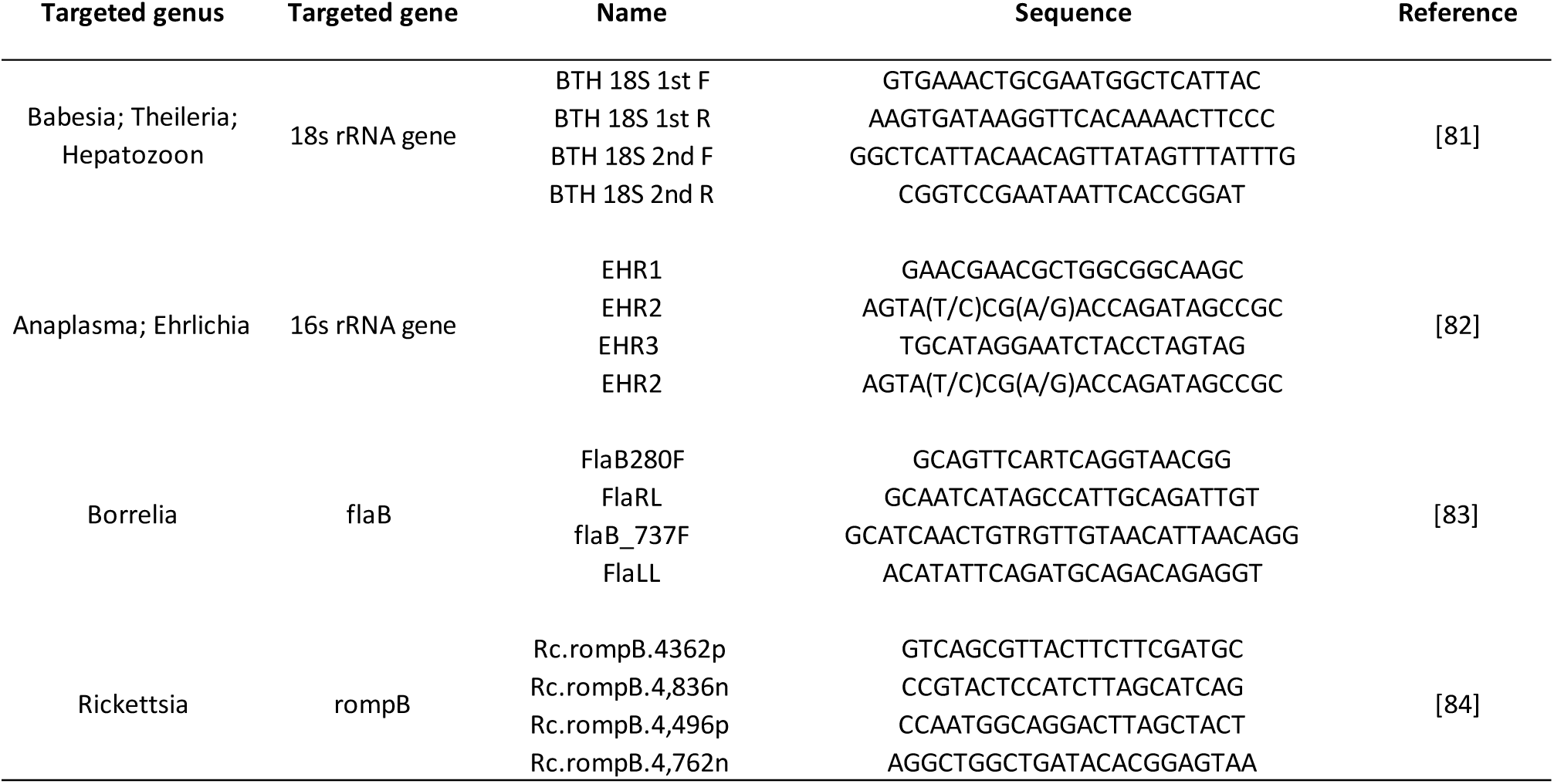

## References

1. Jongejan F, Uilenberg G. The global importance of ticks. Parasitology. 2004;129:S3–14.

2. de la Fuente J, Estrada-Pena A, Venzal JM, Kocan KM, Sonenshine DE. Overview: ticks as vectors of pathogens that cause disease in humans and animals. Front Biosci. 2008;13:6938–6946.

3. Dantas-Torres F, Chomel BB, Otranto D. Ticks and tick-borne diseases: a One Health perspective. Trends Parasitol. 2012;28:437–46.

4. Strle F. Human granulocytic ehrlichiosis in Europe. Int J Med Microbiol Suppl. 2004;293:27–35.

5. Bonnet S, Jouglin M, L’Hostis M, Chauvin A. Babesia sp. EU1 from Roe Deer and Transmission within Ixodes ricinus. Emerg Infect Dis. 2007;13:1208–10.

6. Bonnet S, Jouglin M, Malandrin L, Becker C, Agoulon A, L’hostis M, et al. Transstadial and transovarial persistence of Babesia divergens DNA in Ixodes ricinus ticks fed on infected blood in a new skin-feeding technique. Parasitology. 2007;134:197–207.

7. Cotté V, Bonnet S, Le Rhun D, Le Naour E, Chauvin A, Boulouis H-J, et al. Transmission of Bartonella henselae by Ixodes ricinus. Emerg Infect Dis. 2008;14:1074–80.

8. Bonnet S, Brisseau N, Hermouet A, Jouglin M, Chauvin A. Experimental in vitro transmission of Babesia sp. (EU1) by Ixodes ricinus. Vet Res. 2009;40:1–8.

9. Sprong H, Wielinga PR, Fonville M, Reusken C, Brandenburg AH, Borgsteede F, et al. Ixodes ricinus ticks are reservoir hosts for Rickettsia helvetica and potentially carry flea-borne Rickettsia species. Parasit Vectors. 2009;2:41.

10. Gassner F, van Vliet AJH, Burgers SLGE, Jacobs F, Verbaarschot P, Hovius EKE, et al. Geographic and Temporal Variations in Population Dynamics of Ixodes ricinus and Associated Borrelia Infections in The Netherlands. Vector-Borne Zoonotic Dis. 2010;11:523–32.

11. Reye AL, Hübschen JM, Sausy A, Muller CP. Prevalence and Seasonality of Tick-Borne Pathogens in Questing Ixodes ricinus Ticks from Luxembourg. Appl Environ Microbiol. 2010;76:2923–31.

12. Coipan EC, Jahfari S, Fonville M, Maassen C, van der Giessen J, Takken W, et al. Spatiotemporal dynamics of emerging pathogens in questing Ixodes ricinus. Front Cell Infect Microbiol [Internet]. 2013 [cited 2018 Jul 25];3. Available from: https://www.frontiersin.org/articles/10.3389/fcimb.2013.00036/full

13. Takken W, van Vliet AJH, Verhulst NO, Jacobs FHH, Gassner F, Hartemink N, et al. Acarological Risk of Borrelia burgdorferi Sensu Lato Infections Across Space and Time in The Netherlands. Vector-Borne Zoonotic Dis. 2016;17:99–107.

14. Chvostáč M, Špitalská E, Václav R, Vaculová T, Minichová L, Derdáková M. Seasonal Patterns in the Prevalence and Diversity of Tick-Borne Borrelia burgdorferi Sensu Lato, Anaplasma phagocytophilum and Rickettsia spp. in an Urban Temperate Forest in South Western Slovakia. Int J Environ Res Public Health. 2018;15:994.

15. Gilbert L, Maffey GL, Ramsay SL, Hester AJ. The effect of deer management on the abundance of Ixodes ricinus in Scotland. Ecol Appl. 2012;22:658–67.

16. van Wieren SE, Hofmeester TR. 6. The role of large herbivores in Ixodes ricinus and Borrelia burgdorferi s.l. dynamics. Ecol Prev Lyme Borreliosis [Internet]. Wageningen Academic Publishers; 2016 [cited 2018 Nov 19]. p. 75–89. Available from: https://www.wageningenacademic.com/doi/abs/10.3920/978-90-8686-838-4_6

17. Herrmann C, Gern L. Survival of Ixodes ricinus (Acari: Ixodidae) Under Challenging Conditions of Temperature and Humidity is Influenced by Borrelia burgdorferi sensu lato Infection. J Med Entomol. 2010;47:1196–204.

18. Herrmann C, Gern L. Do the level of energy reserves, hydration status and Borrelia infection influence walking by Ixodes ricinus (Acari: Ixodidae) ticks? Parasitology. 2012;139:330–7.

19. Neelakanta G, Sultana H, Fish D, Anderson JF, Fikrig E. Anaplasma phagocytophilum induces Ixodes scapularis ticks to express an antifreeze glycoprotein gene that enhances their survival in the cold. J Clin Invest. 2010;120:3179–90.

20. Michelet L, Delannoy S, Devillers E, Umhang G, Aspan A, Juremalm M, et al. High-throughput screening of tick-borne pathogens in Europe. Front Cell Infect Microbiol. 2014;4:103.

21. Moutailler S, Moro CV, Vaumourin E, Michelet L, Tran FH, Devillers E, et al. Co-infection of Ticks: The Rule Rather Than the Exception. PLoS Negl Trop Dis. 2016;10:e0004539.

22. Gondard M, Delannoy S, Pinarello V, Aprelon R, Devillers E, Galon C, et al. Upscaling surveillance of tick-borne pathogens in the French Caribbean islands. bioRxiv. 2019;532457.

23. McCullagh P, Nelder JA. Generalized Linear Models. [Internet]. 2nd Edition. London: Chapman et Hall; 1989 [cited 2019 Feb 28]. Available from: https://www.crcpress.com/Generalized-Linear-Models/McCullagh-Nelder/p/book/9780412317606

24. R Core Team. R: A Language and Environment for Statistical Computing [Internet]. Vienna, Austria: R Foundation for Statistical Computing; 2018. Available from: https://www.R-project.org/

25. Vaumourin E, Vourc’h G, Telfer S, Lambin X, Salih D, Seitzer U, et al. To be or not to be associated: power study of four statistical modeling approaches to identify parasite associations in cross-sectional studies. Front Cell Infect Microbiol. 2014;4:62.

26. Malandrin L, Jouglin M, Sun Y, Brisseau N, Chauvin A. Redescription of Babesia capreoli (Enigk and Friedhoff, 1962) from roe deer (Capreolus capreolus): Isolation, cultivation, host specificity, molecular characterisation and differentiation from Babesia divergens. Int J Parasitol. 2010;40:277–84.

27. Dautel H, Dippel C, Kämmer D, Werkhausen A, Kahl O. Winter activity of Ixodes ricinus in a Berlin forest. Int J Med Microbiol. 2008;298:50–4.

28. Perret J-L, Guigoz E, Rais O, Gern L. Influence of saturation deficit and temperature on Ixodes ricinus tick questing activity in a Lyme borreliosis-endemic area (Switzerland). Parasitol Res. 2000;86:554–7.

29. Gilbert L. Altitudinal patterns of tick and host abundance: a potential role for climate change in regulating tick-borne diseases? Oecologia. 2010;162:217–25.

30. Tagliapietra V, Rosà R, Arnoldi D, Cagnacci F, Capelli G, Montarsi F, et al. Saturation deficit and deer density affect questing activity and local abundance of Ixodes ricinus (Acari, Ixodidae) in Italy. Vet Parasitol. 2011;183:114–24.

31. Schulz M, Mahling Monia, Pfister Kurt. Abundance and seasonal activity of questing Ixodes ricinus ticks in their natural habitats in southern Germany in 2011. J Vector Ecol. 2014;39:56–65.

32. Vourc’h G, Abrial D, Bord S, Jacquot M, Masséglia S, Poux V, et al. Mapping human risk of infection with Borrelia burgdorferi sensu lato, the agent of Lyme borreliosis, in a peri-urban forest in France. Ticks Tick-Borne Dis. 2016;7:644–52.

33. Marchant A, Coupanec AL, Joly C, Perthame E, Sertour N, Garnier M, et al. Infection of Ixodes ricinus by Borrelia burgdorferi sensu lato in peri-urban forests of France. PLOS ONE. 2017;12:e0183543.

34. Capelli G, Ravagnan S, Montarsi F, Ciocchetta S, Cazzin S, Porcellato E, et al. Occurrence and identification of risk areas of Ixodes ricinus-borne pathogens: a cost-effectiveness analysis in north-eastern Italy. Parasit Vectors. 2012;5:61.

35. Overzier E, Pfister K, Herb I, Mahling M, Böck G, Silaghi C. Detection of tick-borne pathogens in roe deer (Capreolus capreolus), in questing ticks (Ixodes ricinus), and in ticks infesting roe deer in southern Germany. Ticks Tick-Borne Dis. 2013;4:320–8.

36. Pangrácová L, Derdáková M, Pekárik L, Hviščová I, Víchová B, Stanko M, et al. Ixodes ricinus abundance and its infection with the tick-borne pathogens in urban and suburban areas of Eastern Slovakia. Parasit Vectors. 2013;6:238.

37. Reye AL, Stegniy V, Mishaeva NP, Velhin S, Hübschen JM, Ignatyev G, et al. Prevalence of Tick-Borne Pathogens in Ixodes ricinus and Dermacentor reticulatus Ticks from Different Geographical Locations in Belarus. PLOS ONE. 2013;8:e54476.

38. Hansford KM, Fonville M, Jahfari S, Sprong H, Medlock JM. Borrelia miyamotoi in host-seeking Ixodes ricinus ticks in England. Epidemiol Infect. 2015;143:1079–87.

39. Paul REL, Cote M, Le Naour E, Bonnet SI. Environmental factors influencing tick densities over seven years in a French suburban forest. Parasit Vectors. 2016;9:309.

40. Sormunen JJ, Penttinen R, Klemola T, Hänninen J, Vuorinen I, Laaksonen M, et al. Tick-borne bacterial pathogens in southwestern Finland. Parasit Vectors. 2016;9:168.

41. Raileanu C, Moutailler S, Pavel I, Porea D, Mihalca AD, Savuta G, et al. Borrelia Diversity and Co-infection with Other Tick Borne Pathogens in Ticks. Front Cell Infect Microbiol [Internet]. 2017 [cited 2019 Jan 14];7. Available from: http://journal.frontiersin.org/article/10.3389/fcimb.2017.00036/full

42. Reis C, Cote M, Paul REL, Bonnet S. Questing Ticks in Suburban Forest Are Infected by at Least Six Tick-Borne Pathogens. Vector-Borne Zoonotic Dis. 2010;11:907–16.

43. Marsot M, Henry P-Y, Vourc’h G, Gasqui P, Ferquel E, Laignel J, et al. Which forest bird species are the main hosts of the tick, Ixodes ricinus, the vector of Borrelia burgdorferi sensu lato, during the breeding season? Int J Parasitol. 2012;42:781–8.

44. Marsot M, Chapuis J-L, Gasqui P, Dozières A, Masséglia S, Pisanu B, et al. Introduced Siberian Chipmunks (Tamias sibiricus barberi) Contribute More to Lyme Borreliosis Risk than Native Reservoir Rodents. PLOS ONE. 2013;8:e55377.

45. Reif KE, Macaluso KR. Ecology of Rickettsia felis: A Review. J Med Entomol. 2009;46:723–36.

46. Brown LD, Macaluso KR. Rickettsia felis, an Emerging Flea-Borne Rickettsiosis. Curr Trop Med Rep. 2016;3:27–39.

47. Oliveira K, S Oliveira L, C A Dias C, Silva A, R Almeida M, Almada G, et al. Molecular identification of Rickettsia felis in ticks and fleas from an endemic area for Brazilian Spotted Fever. 2008.

48. Vayssier-Taussat M, Moutailler S, Michelet L, Devillers E, Bonnet S, Cheval J, et al. Next Generation Sequencing Uncovers Unexpected Bacterial Pathogens in Ticks in Western Europe. PLOS ONE. 2013;8:e81439.

49. Lejal E, Moutailler S, Šimo L, Vayssier-Taussat M, Pollet T. Tick-borne pathogen detection in midgut and salivary glands of adult Ixodes ricinus. Parasit Vectors. 2019;12:152.

50. Sprong H, Fonville M, Docters van Leeuwen A, Devillers E, Ibañez-Justicia A, Stroo A, et al. Detection of pathogens in Dermacentor reticulatus in northwestern Europe: evaluation of a high-throughput array. Heliyon. 2019;5:e01270.

51. Pfäffle M, Littwin N, Muders SV, Petney TN. The ecology of tick-borne diseases. Int J Parasitol. 2013;43:1059–77.

52. Herrmann C, Voordouw MJ, Gern L. Ixodes ricinus ticks infected with the causative agent of Lyme disease, Borrelia burgdorferi sensu lato, have higher energy reserves. Int J Parasitol. 2013;43:477–83.

53. Halos L, Jamal T, Maillard R, Beugnet F, Menach AL, Boulouis H-J, et al. Evidence of Bartonella sp. in questing adult and nymphal Ixodes ricinus ticks from France and co-infection with Borrelia burgdorferi sensu lato and Babesia sp. Vet Res. 2005;36:79–87.

54. Schicht S, Junge S, Schnieder T, Strube C. Prevalence of Anaplasma phagocytophilum and Coinfection with Borrelia burgdorferi Sensu Lato in the Hard Tick Ixodes ricinus in the City of Hanover (Germany). Vector-Borne Zoonotic Dis. 2011;11:1595–7.

55. Andersson M, Bartkova S, Lindestad O, Råberg L. Co-Infection with ‘Candidatus Neoehrlichia mikurensis’ and Borrelia afzelii in Ixodes ricinus Ticks in Southern Sweden. Vector-Borne Zoonotic Dis. 2013;13:438–42.

56. Cosson J-F, Michelet L, Chotte J, Le Naour E, Cote M, Devillers E, et al. Genetic characterization of the human relapsing fever spirochete Borrelia miyamotoi in vectors and animal reservoirs of Lyme disease spirochetes in France. Parasit Vectors. 2014;7:233.

57. Castro LR, Gabrielli S, Iori A, Cancrini G. Molecular detection of Rickettsia, Borrelia, and Babesia species in Ixodes ricinus sampled in northeastern, central, and insular areas of Italy. Exp Appl Acarol. 2015;66:443–52.

58. Tijsse-Klasen E, Sprong H, Pandak N. Co-infection of Borrelia burgdorferi sensu lato and Rickettsia species in ticks and in an erythema migrans patient. Parasit Vectors. 2013;6:347.

59. Moniuszko A, Dunaj J, Święcicka I, Zambrowski G, Chmielewska-Badora J, Żukiewicz-Sobczak W, et al. Co-infections with Borrelia species, Anaplasma phagocytophilum and Babesia spp. in patients with tick-borne encephalitis. Eur J Clin Microbiol Infect Dis. 2014;33:1835–41.

60. Hoversten K, Bartlett MA. Diagnosis of a tick-borne coinfection in a patient with persistent symptoms following treatment for Lyme disease. Case Rep. 2018;2018:bcr-2018–225342.

61. Krause PJ, Telford SR, Spielman A, Sikand V, Ryan R, Christianson D, et al. Concurrent Lyme Disease and Babesiosis: Evidence for Increased Severity and Duration of Illness. JAMA. 1996;275:1657–60.

62. Diuk-Wasser MA, Vannier E, Krause PJ. Coinfection by Ixodes Tick-Borne Pathogens: Ecological, Epidemiological, and Clinical Consequences. Trends Parasitol. 2016;32:30–42.

63. Aivelo T, Norberg A, Tschirren B. Human pathogen co-occurrence in Ixodes ricinus ticks: effects of landscape topography, climatic factors and microbiota interactions. bioRxiv. 2019;559245.

64. Strnad M, Hönig V, Růžek D, Grubhoffer L, Rego ROM. Europe-Wide Meta-Analysis of Borrelia burgdorferi Sensu Lato Prevalence in Questing Ixodes ricinus Ticks. Stabb EV, editor. Appl Environ Microbiol. 2017;83:e00609–17.

65. Herrmann C, Gern L, Voordouw MJ. Species co-occurrence patterns among Lyme borreliosis pathogens in the tick vector Ixodes ricinus. Appl Environ Microbiol. 2013;AEM.02158–13.

66. Humair P-F, Postic D, Wallich R, Gern L. An Avian Reservoir (Turdus merula) of the Lyme Borreliosis Spirochetes. Zentralblatt Für Bakteriol. 1998;287:521–38.

67. Kurtenbach K, Carey D, Hoodless AN, Nuttall PA, Randolph SE. Competence of Pheasants as Reservoirs for Lyme Disease Spirochetes. J Med Entomol. 1998;35:77–81.

68. Huegli D, Hu CM, Humair P-F, Wilske B, Gern L. Apodemus Species Mice Are Reservoir Hosts of Borrelia garinii OspA Serotype 4 in Switzerland. J Clin Microbiol. 2002;40:4735–7.

69. Richter D, Schlee DB, Allgower R, Matuschka F-R. Relationships of a Novel Lyme Disease Spirochete, Borrelia spielmani sp. nov., with Its Hosts in Central Europe. Appl Environ Microbiol. 2004;70:6414–9.

70. Richter D, Schlee DB, Matuschka F-R. Reservoir Competence of Various Rodents for the Lyme Disease Spirochete Borrelia spielmanii. Appl Environ Microbiol. 2011;77:3565–70.

71. Taragel’ova V, Koci J, Hanincova K, Kurtenbach K, Derdakova M, Ogden NH, et al. Blackbirds and Song Thrushes Constitute a Key Reservoir of Borrelia garinii, the Causative Agent of Borreliosis in Central Europe. Appl Environ Microbiol. 2008;74:1289–93.

72. Randolph SE, Gern L, Nuttall PA. Co-feeding ticks: Epidemiological significance for tick-borne pathogen transmission. Parasitol Today. 1996;12:472–9.

73. van Duijvendijk G, Coipan C, Wagemakers A, Fonville M, Ersöz J, Oei A, et al. Larvae of Ixodes ricinus transmit Borrelia afzelii and B. miyamotoi to vertebrate hosts. Parasit Vectors [Internet]. 2016 [cited 2019 Jan 14];9. Available from: http://www.parasitesandvectors.com/content/9/1/97

74. Skuballa J, Oehme R, Hartelt K, Petney T, Bücher T, Kimmig P, et al. European Hedgehogs as Hosts for Borrelia spp., Germany. Emerg Infect Dis. 2007;13:952–3.

75. Skuballa J, Petney T, Pfäffle M, Oehme R, Hartelt K, Fingerle V, et al. Occurrence of different Borrelia burgdorferi sensu lato genospecies including B. afzelii, B. bavariensis, and B. spielmanii in hedgehogs (Erinaceus spp.) in Europe. Ticks Tick-Borne Dis. 2012;3:8–13.

76. Gern L, Rouvinez E, Toutoungi LN, Godfroid E. Transmission cycles of Borrelia burgdorferi sensu lato involving Ixodes ricinus and/or I. hexagonus ticks and the European hedgehog, Erinaceus europaeus, in suburban and urban areas in Switzerland. Folia Parasitol (Praha). 1997;44:309–14.

77. Margos G, Vollmer SA, Cornet M, Garnier M, Fingerle V, Wilske B, et al. A New Borrelia Species Defined by Multilocus Sequence Analysis of Housekeeping Genes. Appl Environ Microbiol. 2009;75:5410–6.

78. Hanincova K, Taragelova V, Koci J, Schafer SM, Hails R, Ullmann AJ, et al. Association of Borrelia garinii and B. valaisiana with Songbirds in Slovakia. Appl Environ Microbiol. 2003;69:2825–30.

79. Rauter C, Hartung T. Prevalence of Borrelia burgdorferi Sensu Lato Genospecies in Ixodes ricinus Ticks in Europe: a Metaanalysis. Appl Environ Microbiol. 2005;71:7203–16.

80. Lommano E, Bertaiola L, Dupasquier C, Gern L. Infections and co-infections of questing Ixodes ricinus ticks by emerging zoonotic pathogens in Western Switzerland. Appl Environ Microbiol. 2012;AEM.07961–11.

